# Functional Regrowth of Norepinephrine Axons in the Adult Mouse Brain Following Injury

**DOI:** 10.1101/2024.08.19.608684

**Authors:** Patrick Cooke, David J. Linden

## Abstract

It is widely believed that axons in the central nervous system of adult mammals do not regrow following injury. This failure is thought, at least in part, to underlie the limited recovery of function following injury to the brain or spinal cord. Some studies of fixed tissue have suggested that, counter to dogma, norepinephrine (NE) axons regrow following brain injury. Here, we have used in vivo two-photon microscopy in layer 1 of the primary somatosensory cortex in transgenic mice harboring a fluorophore selectively expressed in NE neurons. This protocol allowed us to explore the dynamic nature of NE axons following injury with the selective NE axon toxin N-(2-chloroethyl)-N-ethyl-2-bromobenzylamine (DSP4). Following DSP4 treatment, NE axons were massively depleted and then slowly and partially recovered their density over a period of weeks. This regrowth was dominated by new axons entering the imaged volume. There was almost no contribution from local sprouting from spared NE axons. Regrown axons did not appear to use either the paths of previously lesioned NE axons nor NE axons that were spared and survived DSP4 treatment as a guide. To measure NE release, GCaMP8s was selectively expressed in neocortical astrocytes and startle-evoked, NE receptor-mediated Ca^2+^ transients were measured. These Ca^2+^ transients were abolished soon after DSP4 lesion but returned to pre-lesion values after 3-5 weeks, roughly coincident with NE axon regrowth, suggesting that the regrown NE axons are competent to release NE in response to a physiological stimulus in the awake mouse.

**Significance Statement:** It is widely believed that axons in the central nervous system (CNS) of adult mammals are incapable of regrowth following injury. Counter to this notion, we describe the structural and functional regrowth of norepinephrine axons following brain injury in the adult mouse. These results extend previous studies describing the regenerative capacity of serotonin axons in the CNS by demonstrating axon regrowth of another neuronal subtype and the capacity of these regrown axons to respond normally to an external physiological stimulus. Taken together, these findings suggest that monoaminergic neurons share a common program for axon regrowth. Elucidation of this molecular and genetic program could inform therapies to promote axon regrowth and functional recovery following injury to the CNS.

## Introduction

It is generally believed that axons within the adult mammalian central nervous system (CNS) are unable to regenerate following injury.^1,2,3^ This lack of axonal regeneration is thought to impede the functional recovery that follows mechanical or chemical brain injury, spinal cord injury, and stroke. A portion of the partial recovery that follows these injuries can be attributed to compensatory sprouting from surviving axons, a process which is distinct from axonal regeneration (which involves regrowth originating from the damaged axon itself, see Tuszynski and Steward, 2012; Cooke et al., 2022)^4,5^.

Importantly, this failure of axonal regeneration in the CNS does not hold for axons that contain the monoamine neurotransmitter serotonin. Immunohistochemical studies in damaged spinal cords of adult mice and rats treated with grafts and chondroitinase have shown that certain serotonin axons, which originate from the caudal raphe nuclei, completely crossed the graft and, in so doing, contributed to the restoration of bladder control, diaphragm control and locomotion (see Perrin and Noristani, 2019 and Cooke et al., 2022, for review)^6,7,8,9,10,11,12,5^.

In brain tissue of adult mice and rats, immunohistochemical studies have shown serotonin axon regrowth following amphetamine-induced lesions^13,14,15,16,17,18^, thermal injury,^19^ and traumatic brain injury resulting from a neocortical stab^17^ or controlled cortical impact.^20^ Unlike the spinal cord, the serotonin axon regrowth in these brain injury studies proceeded without any therapeutic intervention.

Immunohistochemical studies are limited by their across-animal design and are thus unable to distinguish sprouting originating from surviving serotonin axons within the field of view from regeneration, which is the growth from the damaged axons themselves. Using long-term in vivo imaging of serotonin axons in the neocortex of serotonin transporter-EGFP BAC transgenic mice, we were able to show new growth from > 80% of the cut ends of serotonin axons in the weeks following neocortical stab injury. Often these regrowing axons were able to completely cross the glial that formed within.^17^ When repeated amphetamine treatment was used to lesion serotonin axons, in vivo imaging revealed that the subsequent slow recovery of axon density in the neocortex was dominated by long-distance regrowth from outside the field of view with little contribution from local sprouting. Furthermore, unlike certain regrowing axons in the peripheral nervous system,^21^ new serotonin axons did not use surviving axons as a guide for regrowth, nor did they follow the pathways left by degenerated axons. These regrown axons then survived at the same rate as serotonin axons in control mice which did not receive amphetamine treatment.

Might the serotonin neuron’s unusual capacity to regrow their axons unaided following brain injury be shared by other monoamine neuromodulators? Several immunohistochemical experiments have suggested that this is the case for norepinephrine (NE) axons. Lesions to the superior peduncle of the adult rat cerebellar cortex resulted in increased arborization of terminal NE-immunopositive fibers.^22^ Following chemical damage with a selective NE, neurotoxin N-(2-chloroethyl)-Nethyl-2-bromobenzylamine, DSP-4,^23,24^ NE immunoreactive axons were first lost and then subsequently recovered by 12 months post-lesion in rats.^25,26^ Finally, a recent study showed reduction of NE axon density following controlled cortical impact or the application of a cortical stab wound 1 week and 1 month following injury in mice. Much like serotonin axons, NE axon density slowly recovered over the course of several months.^27^ As with the initial evidence of serotonin axon regeneration, these fixed tissue studies of NE axons were unable to distinguish regeneration from collateral sprouting.

Here, we used in vivo two-photon imaging of transgenic mice selectively expressing tdTomato in NE neurons to monitor axon regrowth in primary somatosensory cortex over the course of 16 weeks following selective chemical lesioning with DSP4. This method allowed us to measure the dynamics of NE axon regrowth and to test the hypothesis that growth and survival patterns are conserved between regrowing NE and serotonin axons.

Additionally, we sought to test the hypothesis that the post-DSP-4 recovery of NE axon density is accompanied by a recovery in functional NE signaling. To do so, we used a recombinant adeno-associated virus (AAV) to express a Ca^2+^ indicator in astrocytes of the primary somatosensory cortex of transgenic mice bearing labelled NE axons. By using the well-characterized α1-NE-receptor mediated startle response in neocortical astrocytes of awake mice,^28,29^ we employed an index of functional NE release to address this key question. In so doing, we are able to answer the question of whether regrown NE axons are competent to release NE and thereby potentially contribute to the recovery of brain function after injury.

## Results

### Immunohistochemical measurements show the regrowth of NE axons following DSP4- induced degeneration

NE axons originate from somata within the locus coeruleus and project throughout the brain. Those NE axons which extend into the forebrain innervate the thalamus, amygdala, hypothalamus, and nucleus accumbens before turning dorsally and finally posteriorly to innervate the cerebral cortex, creating a C-shape when viewed in the sagittal plane. In the cerebral cortex, these axons primarily run from anterior to posterior in layers 1 and 5 and project collaterals into the intermediary layers 2, 3 and 4 (Figure 1A).^30,31,32,33^

**Figure 1.**
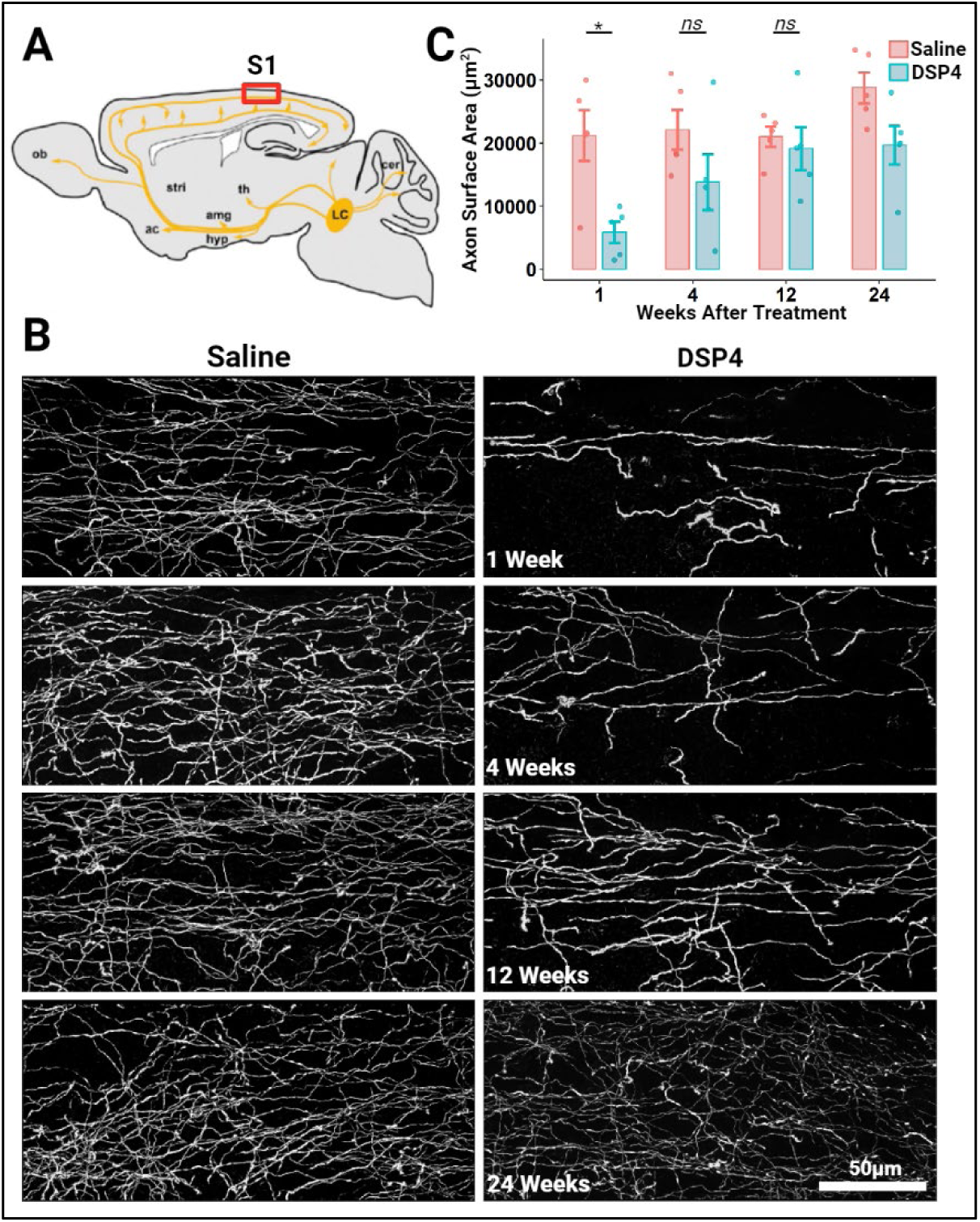
NE axons innervating the adult mouse primary somatosensory cortex regrow following a chemical lesion with the NE-specific neurotoxin DSP4 (50mg/kg). ***A***, Schematic diagram of a sagittal section of an adult mouse brain depicting the general projection of axons from NE cell bodies located within the locus coeruleus (LC). The red box indicates the region where this histological analysis was conducted: the primary somatosensory cortex (S1). *\*Other abbreviations: olfactory bulb (OB), striatum (stri), nucleus accumbens (ac), amygdala (amg), hypothalamus (hyp), thalamus (th), cerebellum (cer). **B***, Representative maximum projected confocal stack images of layer 1 of the primary somatosensory cortex in mice sacrificed 1-, 4-, 12-, and 24-weeks following treatment and sliced in the sagittal plane. Dopamine-β-hydroxylase (DBH)-cre x mTmG mice were used to selectively label NE axons, neuronal tissue was sliced along the sagittal plane, and the fluorescent signal was amplified through processing with antibodies raised against GFP. ***C***, IMARIS software was used to quantify the total axon surface area within 3-D reconstructed z-stacks (z=30µm) of layer 1. Each plot symbol represents the total axon surface area of a single sagittal section of an individual mouse (n=5/group) and vertical bars show the standard error. * = *P* < 0.05; *ns* = not significant.

To test the hypothesis that NE neurons are capable of regrowing their axons following injury, we first sought to replicate and extend previous studies which demonstrated the rapid loss and subsequent slow return of axons throughout the forebrain immunoreactive for either NE itself or the key NE synthesizing enzyme, dopamine-β-hydroxylase (DBH), following treatment with the selective NE axon neurotoxin, DSP4, in rats.^25,26^ It is formally possible that the loss and slow return of immunoreactivity does not reflect the loss of axons but rather the loss of NE and DBH from intact axons. To address this possibility, we utilized DBH-cre mice crossed with the mTmG reporter line^34^ to generate mice that selectively and continually express membrane-bound EGFP within NE neurons under the control of a strong pCA promoter (DBHcre x mTmG). Previous work from our lab has shown that these EGFP-immunoreactive axons in layer 1 are immunopositive for tyrosine hydroxylase, the enzyme that synthesizes both NE and dopamine, but immunonegative for the dopamine transporter, consistent with them being noradrenergic.^27^

DBHcre x mTmG mice between the ages of 15 and 25 weeks were separated by age and sex before being randomly assigned to receive either a single 50 mg/kg systemic dose of DSP4 or an equal volume of saline. At each of four time points −1, 4, 12, and 24 weeks following treatment administration −5 age and sex matched mice were sacrificed and their brains were fixed and sliced in the sagittal plane. Slices were then immunolabelled with antibodies raised against EGFP in order to amplify the EGFP fluorescence within the membrane of NE axons. Images of layers 1, 2/3, and 5 reveal the dramatic loss and subsequent recovery of NE axons following treatment with DSP4 in comparison to saline treated mice across neocortical layers in both primary somatosensory (Figures 1B, S1A) and motor (Figure S2A) cortex. Measurement of the total surface area of NE axons in layer 1 of the primary somatosensory cortex one week following DSP4 treatment (5844 ± 1688 μm^2^, mean ± SE) revealed a significant (*P* = 0.015) decrease compared to those treated with saline (21206μm^2^ ± 4013μm^2^). 12 weeks later, total NE axon surface area in DSP4-treated animals had recovered significantly (19127μm^2^ ± 3402μm^2^, *P* = 0.019) and was no longer significantly different from saline-controls (20980μm^2^ ± 2448μm^2^, *P* = 0.641) (Figure 1C). Similar changes were seen across cortical layers 1 - 5 in both the somatosensory and motor cortices (Figure S1B, S2B). These results demonstrate that NE axons slowly regrow unaided following DSP4-mediated injury. However, the cross-animal design inherent to immunohistochemical experiments prevents us from determining the extent to which this return is due to the collateral sprouting from surviving axons or regeneration originating from the shaft or the ends of damaged axons.

### Regrowth of NE axons is dominated by new growth and not local sprouting from surviving axons

To measure the dynamics of axonal regrowth, we employed in vivo two-photon microscopy through a cranial window overlying the primary somatosensory cortex. For these experiments, we crossed DBH-cre mice with the Ai14 reporter line to generate mice that selectively express the soluble red tdTomato fluorophore within NE neurons (DBH-cre x Ai14) (Figure S3). Changing the reporter line from mTmG to Ai14 allowed us to use the green channel for simultaneous Ca^2+^ imaging with GCaMP8s. The somatosensory cortex of both male and female DBH-cre x Ai14 mice aged 9-12 weeks were infected with AAV to induce the expression of the fluorescent Ca^2+^ indicator, GCaMP8s, and glass windows were then implanted to seal the craniotomy. After a 6 week recovery period to allow for stable viral expression and the resolution of acute inflammation from the injection and window surgery, an initial z stack encompassing layer 1 was acquired followed by a second pretreatment image of the same region taken 2 weeks later. The following day, mice were treated with either DSP4 (50 mg/kg, n=10) or saline (n=9). Imaging of the same region then continued at two-week intervals until the imaging window clarity deteriorated.

While prior studies examining the effects of DSP4 have established a dose-dependent lesioning effect on NE axons, they show inconsistent results in regards to the effect on NE cell bodies at the 50mg/kg dose. Fritschy and Grzanna^35,26^ demonstrated a pronounced loss of DBH immunopositive cell bodies 6 months following DSP4 treatment in rats. These results were corroborated by Zhang et al. (1995)^36^ who used Nissl-staining to demonstrate the loss of locus coeruleus cell bodies 3 months following DSP4 treatment in rats. In 2010 however, Szot et al. found no DSP4 induced loss of NE cell bodies in rats using *in situ* hybridization for tyrosine hydroxylase, norepinephrine transporter, and DBH. Finally, Iannitelli et al., (2023)^37^ administered 2 50mg/kg doses of DSP4 (given 1 week apart) and saw no NE cell body loss 2 weeks following the final dose as assed through staining for DAPI and immunostaining for tyrosine hydroxylase. These differences are likely due to a variety of factors including the staining method used and the time points sampled after DSP4 administration.^38^

To address this question in our experiments, the same DBHcre x Ai14 mice assessed in Figure 2 were sacrificed 24 weeks following treatment and their brains fixed and sliced in the sagittal plane. Slices were then immunolabelled with antibodies raised against DsRed in order to amplify the td-Tomato fluorescence within NE neurons. Exhaustive imaging and counts of Ai14- positive cell bodies within the locus coeruleus of the right hemisphere showed consistency between groups (DSP4 = 1246 ± 134 somata, n = 9; Saline = 1509 ± 204, n = 8; *P* = 0.289), indicating that treatment with 50mg/kg of DSP4 lesioned NE axons but spared the cell bodies (Figure S4).

**Figure 2.**
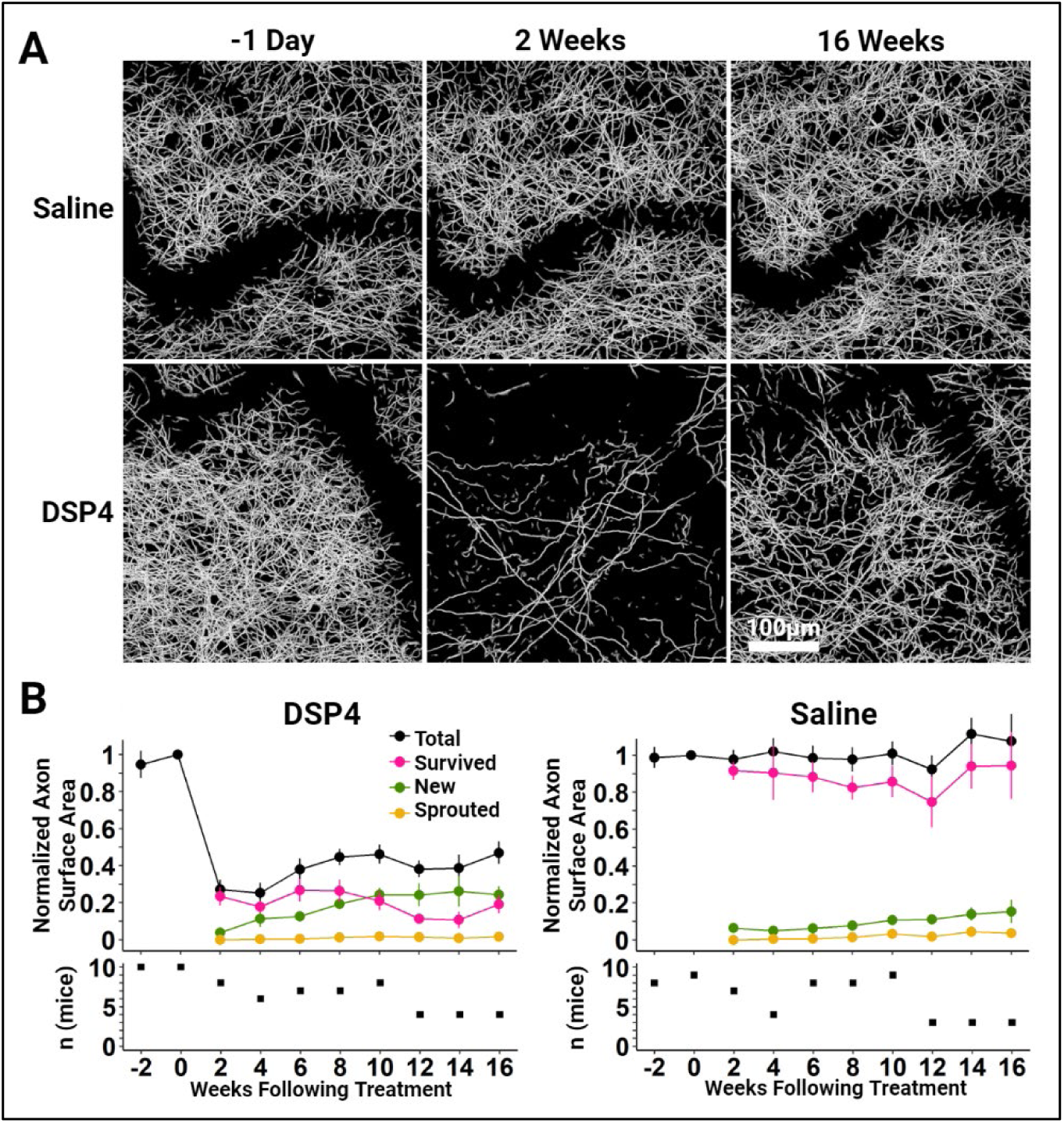
Long-Term *in vivo* 2-photon imaging shows the regrowth of NE axons innervating the adult mouse neocortex after DSP4 challenge. ***A***, Representative 108μm thick maximum projected stack images of layer 1 of the primary somatosensory cortex show the widespread loss and slow recovery of NE density over 16 weeks in a representative DSP4-treated mouse. In a representative saline-treated mouse, the overall axon density and individual axon location and morphology was quite stable. Specific labeling of NE axons was achieved using DBHcre x Ai14 transgenic mice. The dark, sinuous patches in these images are the shadows created by large surface blood vessels. ***B***, (top) Population time-course data measuring total axon surface area are normalized to the value measured 1 day prior to treatment. Recovery of axon density following DSP4 treatment is dominated by new axon growth. There is almost no contribution from collateral sprouting originating from survived axons. Axons that initially survived DSP4-treatment continued to survive at a rate similar to those that were saline-treated. Vertical bars show the standard error. The number of mice imaged at each timepoint, shown in the lower panel, varied due to the deterioration of the imaging window’s clarity over time and downtime due to repair of the imaging rig from week to week. This change in the number of mice assessed at each timepoint, coupled with the variation in lesion magnitude across DSP4-treated animals (SE of Survived axons = 5.0% at week 2), led to some apparent instability across time in the population measures. However, examining survived axons within individual animals shows consistent stability over 16 weeks (Figure S4).

Figure 2A shows representative maximally projected z stack images of layer 1 of the somatosensory cortex obtained 1 day prior to treatment, 2 weeks after treatment, and then 16 weeks after treatment. Calculation of the total surface area of NE axons revealed extensive degeneration in mice treated with DSP4, which decreased to 27.0 ± 5.5% (n = 8; *P* < 0.001) of the pretreatment baseline. Saline-treated animals saw no significant change (97.7 ± 5.2% of baseline; n = 7; *P* = 0.672). Over the subsequent weeks, axon surface area gradually returned to the imaged volume, reaching 46.2 ± 5.2% (n = 8) of the pre-treatment baseline 10 weeks later, a significant improvement from week 2 (*P* = 0.002). Total axon surface area of the saline treated group remained stable over that period (101 ± 6.4% at 10 weeks; n = 9, *P* = 0.465; Figures 2B and S4).

To further analyze this partial recovery of NE axon density following DSP4 treatment, axon surfaces were broken into segments which we assigned to one of several groups: surviving axons, which persisted after treatment, sprouted axons, which grew as branches elaborated from surviving axons following treatment, and new axons, which appeared in the analyzed volume only after treatment. Almost none of the recovery following DSP4 treatment resulted from collateral sprouting, which measured 0.0 ± 0.0%, after 2 weeks, and 1.7 ± 0.3%, after 10 weeks of the total pre-treatment surface area (n = 8 each; *P* < 0.001). Similarly, little collateral sprouting was seen in mice treated with saline (3.3 ± 0.2% at 10 weeks, n = 9, *P* < 0.001). Over this same interval, the surface area of new axons in DSP4-treated mice increased from 3.8 ± 0.7%, after 2 weeks, to 24.4 ± 3.4%, after 10 weeks, of the total-pre-treatment surface area (n = 8 each; *P* < 0.001), far outpacing the changes seen in saline-treated mice which increased from 6.3 ± 1.2%, after 2 weeks, to 10.6 ± 1.0%, after 10 weeks (n = 7 and 9 respectively; *P* < 0.001). Axons that survived DSP4 treatment saw no significant change from week 2 through week 10 following treatment, respectively measuring 23.4 ± 5.0% and 20.9% ± 4.9% of the total pre-treatment surface area (n = 8 each; *P* = 0.270) (Figure 2B and S4). The long-term stability of axons that survived DSP4 treatment was similar to the saline treated group which also saw no significant change, measuring 91.5 ± 5.0% 2 weeks after treatment and 85.7 ± 8.5% at 10 weeks (n = 7 and 9 respectively; *P* = 0.374). The stability of survived axons in both groups not only demonstrates that axons are stable following DSP4-treatment but also that NE axons are stable over the course of months in the absence of perturbation.

Are the new axons that appear in the analyzed volume after DSP4 treatment the result of true regeneration from damaged ends or rather collateral sprouting from survived axons at some location outside of the imaging volume? Although we cannot rule out this latter possibility from the present observations, 16 weeks following DSP4 treatment (n = 4) the average length of all sprouted axons within the imaging volume was 10.8μm ± 6.0μm, the longest of which measured 30.7μm ± 18.7μm. These results were similar in saline-treated animals (n = 3) whose average sprout length measured 8.1μm ± 0.42μm and the longest sprouts were 18.2μm ± 5.1μm. While collateral sprouting from survived axons does occur, it seems to be quite limited in length and so is unlikely to contribute to the extensive regrowth seen in these experiments.

### Regrown NE axons do not appear to use surviving axons as contact guidance cues, nor do they preferentially regrow along the same paths that NE axons occupied prior to treatment

What cues might NE axons use to guide their regrowth from the locus coeruleus through the somatosensory cortex? Could they be using surviving axons as scaffolds to guide their regrowth? To answer this question, we measured the location of new and survived axons 6, 10, and 16 weeks following treatment to examine their trajectory (Figure 3A). 10 weeks following DSP4 treatment, only 13.7 ± 6.4% (n=7) of new axon segments were within 5μm of the nearest survived axon segment, in comparison to 41.7 ± 3.2% (n=7) seen in saline treated animals. A higher proportion of new axon segments in close proximity to survived segments in saline-treated mice, as compared to those treated with DSP4, is to be expected if new axons are not using survived axons as guidance cues, merely due to the much higher density of survived axons in saline treated mice

**Figure 3.**
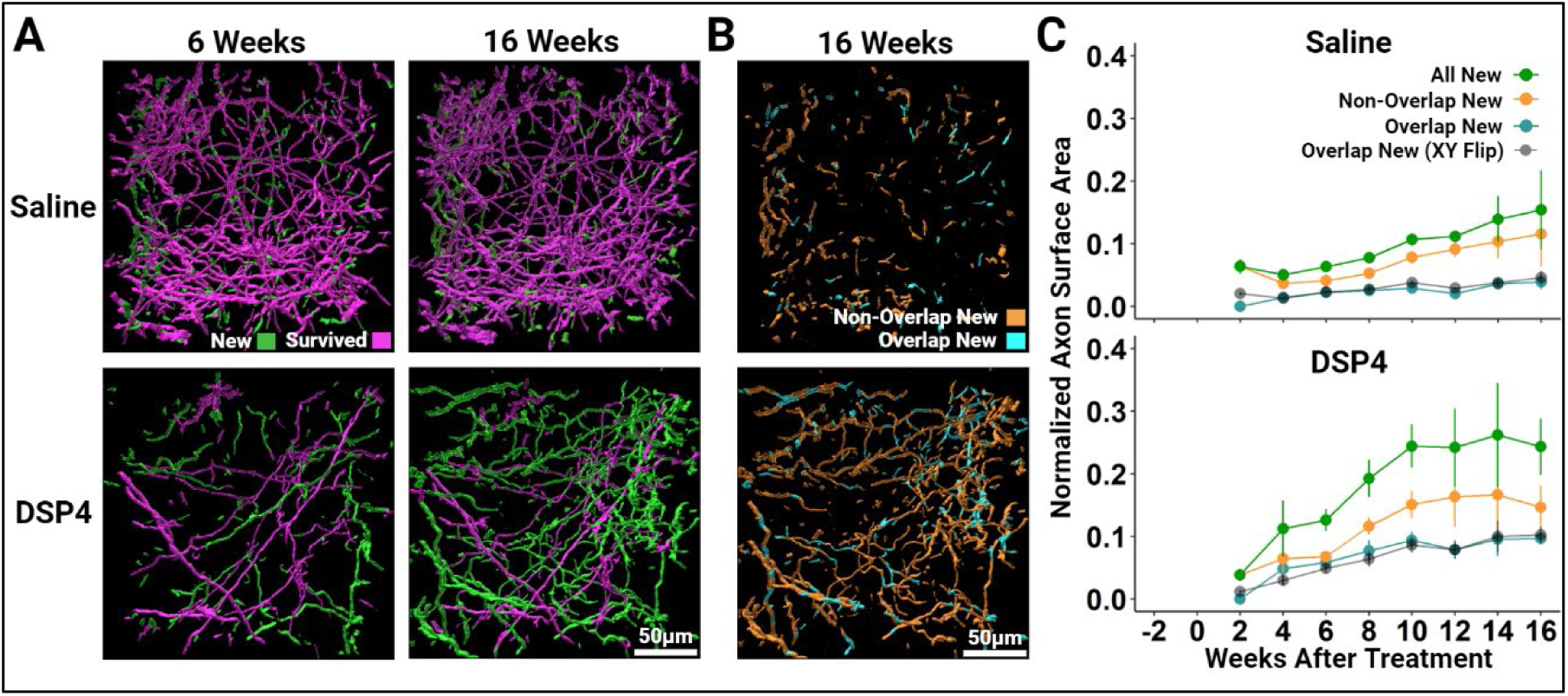
Imaging the regrowth trajectory of NE axons innervating the adult mouse primary somatosensory cortex after DSP4 treatment reveals that they do not appear to be contact-guided by survived axons, nor do they preferentially regrow where axons were present before the DSP4 lesion. ***A***, Representative 3-D images of segmented NE axons innervating the primary somatosensory cortex 6 and 16-weeks following treatment. Two weeks after treatment, axon segments were classified as Survived if they lay within 5µm of the location of an axon segment present 1 day prior to treatment. All other segments are labeled New as they are likely the result of growth originating from outside of the analyzed volume. In all subsequent weeks, segments are categorized as Survived if they lay within 5μm of those classified as Survived at the 2-week timepoint and New if they are 5μm or further. New axons do not appear to be regenerating along Survived axons and thus are unlikely to use surviving axons as a scaffold guiding their regrowth. ***B***, New axon segments, shown in *A,* were isolated and further sub-categorized according to their location relative to the location of axon segments present 1 day prior to injury. New axon segments that lie within 5µm of a segment present 1 day prior to injury are categorized as Overlap new (cyan) while those that lie 5µm or further are categorized as Non-Overlap New (orange). ***C***, Population time-course data measuring the total surface area of segments categorized as All New (green; labelled New in Figure 2B), Non-Overlap New (orange), Overlap New (cyan), and Overlap New (XY flip) (black) are normalized to the total surface area of all segments 1 day prior to treatment. The black line depicts the normalized surface area of axon segments that lie within 5μm of the relative location of axon segments present 1 day following treatment when that pre-treatment volume is flipped on its X and Y axis. This provides an index of the expected proportion of Overlap New axons due to chance crossings. Note that the Overlap New and Overlap New (XY Flip) measurements are very similar, suggesting that nearly all of the Overlap New scoring can be attributed to chance crossings rather than regrowth within the trajectories of previously-lesioned axons. Vertical bars depict the standard error of the mean.

If new axons are not regrowing along survived axons, do they instead grow along paths that had previously been established by NE axons which then degenerated following DSP4 treatment? To investigate this possibility, we isolated new axon segments and then further sub-categorized them based on the location of their center of mass relative to the position of the center of mass of axon segments present 1 day prior to treatment. New axon segments whose center of mass lay within 5µm of the center of mass of an axon segment present prior to treatment were assigned to the ‘Overlap New’ group while those 5µm or further were categorized as ‘Non-Overlap New’ (Figure 3B). 5µm was chosen as a relative positional threshold to allow for small variations in the geometry and orientation of the imaged volume that arise from factors like head placement relative to the imaging plane of the objective lens. Surface area calculations revealed that new axon growth 10 weeks following DSP4 treatment is dominated by axon segments that do not overlap in space with those that were destined to degenerate following DSP4 treatment (Non-Overlap New segments = 15.1 ± 2.2% of the total pretreatment surface area) but with some contribution from Overlap New segments (9.3 ± 1.4% of the total pretreatment surface area, n=8). Saline-treatment revealed comparatively less normalized surface area of both the Non-Overlap New (7.8 ± 1.0%) and Overlap-New (2.9 ± 0.6%) axon segments (n = 9), however it is difficult to directly compare the saline and DSP4 groups in this regard due to the dramatic difference in axon survival following treatment as well as the relative difference in new axon growth.

Even if axons are not regrowing along a guided trajectory, some overlap is expected nonetheless due to chance crossings. To generate an index of the expected proportion of Overlap New axons due to chance crossings, we flipped the analyzed volume 1 day prior to treatment on both its X and Y axis and reanalyzed the extent of overlap. Comparing the area under the curve from weeks 2 through 16 demonstrated that these measurements are quite similar to the Overlap New measurements in both DSP4 (exact *P* = 0.25) and saline (exact *P* = 0.5) treated animals indicating that nearly all of the Overlap New axon segments can be attributed to chance crossings rather than regrowth along the trajectories of previously lesioned axons (Figure 3C).

### Regrown NE axons are capable of releasing neurotransmitter in response to a physiological stimulus

While we have shown that new NE axons regrow through the imaged volume in layer 1 of the somatosensory cortex following DSP4 treatment, the presence of these axons does not demonstrate their ability to release NE in a physiologically relevant manner. To address this functional question, we sought to determine whether these axons are capable of NE release in response to startle. NE neurons in the locus coeruleus fire reliably in response to startle leading to broad release of NE throughout the brain.^39,40^ This NE binds to α1-adrenoreceptors on neocortical astrocytes and elicits a robust Ca^2+^ transient that can be monitored in vivo using fluorescent Ca^2+^ sensors such as GCaMP.^28,29^ Here, in the same DBHcre x Ai14 animals examined in Figures 2 and 3, we infected layer 1 of the somatosensory cortex with an AAV containing a transgene encoding GCaMP8s driven by the astrocyte-selective truncated GFAP promoter, gfaABC1D.^41^ This infection produced patchy GCaMP expression that displayed the tiled patterning, S110β immunoreactivity and filamentous morphology characteristic of astrocytes and was stable over the course of many weeks (Figure S6). To further validate our model, in a separate group of mice we confirmed that increases in GCaMP8s fluorescence evoked by air puff were abolished by treatment with the selective α1-adrenoreceptor antagonist, prazosin (7.5mg/kg, Figure. 4B,C). This strategy allowed us to assess the functional capability of regenerating NE axons by monitoring GCaMP8s fluorescence within astrocytes using startle as an easily manipulated environmental trigger for neurotransmitter release.

**Figure 4.**
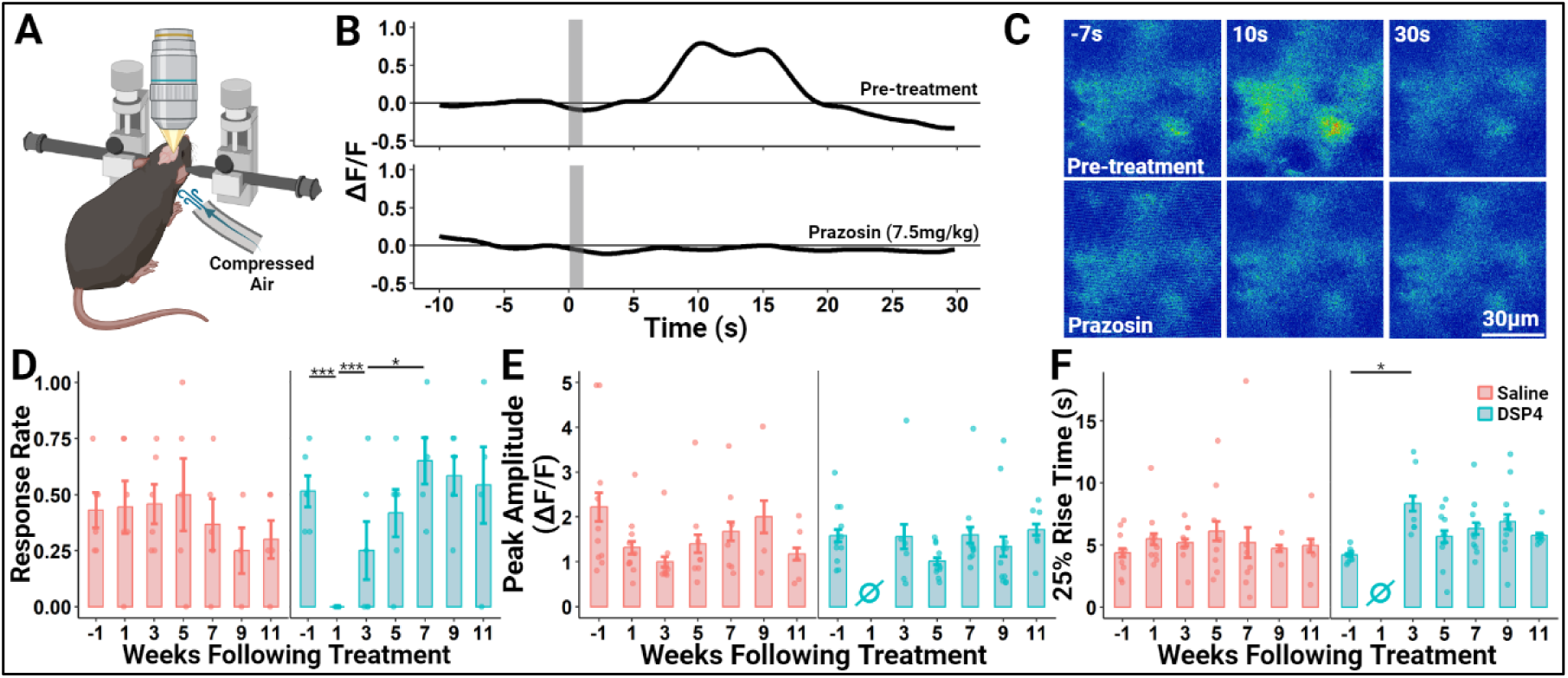
Regrown NE axons are competent to release NE as measured using started-evoked NE-mediated Ca^2+^ transients in astrocytes. ***A***, Schematic of the functional imaging configuration. The primary somatosensory cortex of DBHcre x Ai14 mice was infected with AAV5-gfaABC1D-jGCaMP8s (designed to drive GCaMP8s expression in astrocytes) 5-7 weeks prior to recording. The awake mouse was placed on a treadmill (not depicted) and head fixed. Compressed air (30PSI) was delivered through a tube to the back of the neck to elicit a startle-response and, following a short delay, NE release. ***B***, Representative single trial showing a Ca^2+^ transient measured in a group of adjacent astrocytes evoked by a 1s long air-puff (grey bars) to the back of the mouse’s neck. Treatment with the selective α1-adrenoreceptor antagonist prazosin (7.5mg/kg) 15 minutes prior to the trial abolishes the startle-evoked astrocytic Ca^2+^ transient, indicating that it is mediated by NE release. ***C***, Representative individual (non-averaged) false-color thermal-coded false color images of GCaMP8s fluorescence before and after treatment with 7.5mg/kg Prazosin. ***D-F***, Select response characteristics before and after treatment with the NE-specific neurotoxin DSP4 (50mg/kg) or saline. During each assessment week, animals underwent 4 individual trials with a 5-minute inter-trial interval. Individual trials were excluded from analysis using a baseline stability criterion (see methods). Each plot point represents an individual mouse and vertical bars show the standard error. ***D***, The proportion of trials each assessment week that elicited a startle-evoked Ca^2+^ response in neocortical astrocytes. ***E***, The peak amplitude of all GCaMP8s response traces each week. ***F***, The latency from stimulus onset to 25% of the maximum amplitude of all GCaMP8s response traces each week. * = *P* < 0.05; *** = *P* < 0.001.

After a 6 week recovery period to allow for stable viral GCaMP8s expression, animals were progressively habituated to handling and the imaging rig each day over the course of 7 days and then challenged with an initial series of startle-eliciting air puffs to establish that they do indeed respond with astrocyte Ca^2+^ transients. One week following this initial baseline recording, animals were treated with either 50mg/kg of DSP4 or saline. Beginning 1 week after treatment, air puff-evoked Ca^2+^ transients were measured in those same astrocytes every 2 weeks until the imaging window’s clarity deteriorated. In this way, structural and functional assessments were conducted on alternate weeks to prevent any interference that anesthesia might have on the mouse’s startle response and to reduce habituation to the air puff startle. During each week of functional data collection, alert mice were placed on a treadmill and head fixed to the imaging rig (Figure 4A). Mice were given a period of 15 minutes to rest and adjust to head fixation before undergoing 4 sequential startle challenge trials, each separated by a 5 minute rest period to reduce habituation to the stimulus. Each trial consisted of 40s of continuous recording. Baseline GCaMP8s fluorescence was measured for the first 10 seconds followed by a 1-second-long air puff applied to the back of the mouse’s neck and a 30-second-long period to capture the response.

We examined three features across the 4 GCaMP response trials recorded each week of data collection: the response rate across each of the 4 trials, peak ΔF/F amplitude within each trial, and the time from stimulus onset to 25% of the peak ΔF/F amplitude. This ‘25% rise time’ serves as a measure for latency to the onset of the response. Response rate was calculated as the proportion of the total number of trials that elicited an increase in GCaMP fluorescence intensity, defined as a peak amplitude ΔF/F > 0.5 and a peak z-score > 6. Some trials were excluded from assessment due to fluorescent signal instability during the initial baseline recording (see methods for criteria). Application of the air puff stimulus elicited a response rate of 51.4 ± 6.9% 1 week prior to DSP4 treatment (n = 6). 1 week following treatment this response rate dropped to 0 ± 0% (n = 6) (*P* = 0.000). Over the subsequent weeks, this response rate gradually returned, showing significant improvement by 3 weeks following treatment (25 ± 12.9%, n = 6, *P* = 0.000). By 7 weeks following treatment the response rate had further increased beyond that seen at 3 weeks to 65 ± 10.3% (n = 5, *P* = 0.032). Meanwhile, the response rate in saline treated animals remained stable, demonstrating no significant changes across this period (pre-treatment = 43.1 ± 7.9%, n = 6; week 1 = 44.4 ± 11.2%, n = 6; week 3 = 45.8 ± 8.9%; week 7 = 36.7 ± 11.4%, n = 5) (Figure 4D).

When examining changes in peak amplitude and response latency, it is important to note that these metrics can only be analyzed in trials during which a response was elicited by the stimulus. For example, one week following DSP4 treatment, application of an air puff stimulus elicited no responses and thus there was no peak amplitude or 25% rise time data during that week. While DSP4 treatment did not significantly affect the peak amplitude of the GCaMP response (Figure 4E), there was a significant increase in the 25% rise time 3 weeks following DSP4 treatment in comparison to the pre-treatment latency, which increased from 4.2 ± 0.1s to 8.3 ± 0.6s (n = 11 and 6 respectively; *P* = 0.036). No significant change in 25% rise time was observed in saline treated animals (week 1 = 4.4 ± 0.3s, n = 10; week 3 = 5.2 ± 0.3s, n = 10). These results demonstrate that DSP4 treatment eliminates air puff elicited Ca^2+^ responses in cortical astrocytes 1 week following treatment which gradually return over the course of 7 weeks and that the latency of onset for these responses is increased during the earliest stages of this return.

## Discussion

We investigated the capacity of NE axons to regrow following injury with the selective NE neurotoxin, DSP4 in adult mice. Previous immunohistochemical work in adult rats^25,26^ has described the loss and subsequent return of NE and DBH immunoreactivity following treatment with DSP4 and has suggested that DSP4 treatment induced the selective degeneration of NE axons which then slowly regrew over the following weeks. While this interpretation is likely to be correct, it is formally possible that these findings instead resulted from a DSP4-evoked loss of DBH and NE and subsequent slow anterograde refilling of intact NE axons.

Here, we used DBH-Cre x mTmG mice to selectively label NE neurons with membrane-bound EGFP. In these mice, EGFP is continually produced under the control of a strong viral promoter and hence is unlikely to be rapidly down-regulated following DSP4 treatment. We observed widespread loss of EGFP-positive axons which then slowly regrew over the course of weeks across all layers of primary somatosensory (Figures 1 and S1) and motor (Figure S2) cortices.

While these immunohistochemical experiments confirmed and extended the previous work using NE and DBH immunolabeling in rats,^25,26^ they also relied on cross-animal comparisons of fixed tissue and hence were unable to measure the dynamics of this regrowth or the functional status of NE release from the regrown axons. To address these questions, we employed in vivo two-photon microscopy to monitor axons within individual animals throughout this regrowth period. We again used the DBH-cre system to fluorescently label NE neurons, this time employing the Ai14 reporter mouse line which yields selective expression of the soluble red tdTomato fluorophore (Figure S3). The superficial layers of the somatosensory cortex of these mice were then infected with an AAV engineered to drive expression of GCaMP8s in astrocytes (Figure S6). This combination allowed us to measure the structural regrowth of NE axons in red and astrocyte Ca^2+^ transients in green as an index of local NE release.

In vivo imaging experiments revealed that, following DSP4 lesioning, the return of NE axon density was dominated by new growth from outside of the imaged volume with very little contribution from collateral sprouting (Figures 2 and S5). Furthermore, regrowing axons did not appear to use surviving axons as a guide (Figure 3A) nor did they regrow along the same paths that had been previously established by the degenerated fibers (Figure 3B, C). While it is possible that this growth following DSP4 treatment is the result of collateral sprouting that originates at locations outside of the imaged volume, we believe this to be unlikely as the average length of sprouted collaterals within our imaged volume 16 weeks following treatment was only 10.8μm ± 6.0μm (n = 4). The dynamics of NE axon regrowth closely resemble those of serotonin axons following lesioning with the selective serotonin neurotoxin parachloroamphetamine (PCA). Serotonin axons also showed robust regrowth with little contribution from collateral sprouting. Similarly, they did not appear to use spared axons as a guide or regrow along paths that had been previously established by the degenerated fibers.^17^

While the regrowth of NE axons is clear, determining whether those axons are competent to release NE is not as straight forward. Neurotransmitters such as glutamate, GABA, and glycine are released into classical synapses where their action on ionotropic receptors can be directly measured as current flow across the post-synaptic membrane. However, NE signals through metabotropic receptors, which do not evoke rapid transmembrane currents. Here, we measured release of NE by monitoring α1-adrenoreceptor-mediated internal Ca^2+^ transients in neocortical astrocytes as evoked by startle.^28,29^ These transients were completely eliminated 1 week following DSP4 but gradually returned, beginning 3 weeks following treatment.

While these results provide evidence of a functional response to startle-induced NE release, this method has its limitations. In our experience, startle-evoked Ca^2+^ transients, even in control conditions, were rather variable. NE-evoked Ca^2+^ fluctuations in neocortical astrocytes are driven through mobilization from internal stores which may have variable filling status. Also, the recording room was not entirely soundproof and noises from outside sometimes evoked spontaneous transients which might produce habituation to the startle. Thus, quantification of amplitude or kinetics of the air puff-evoked Ca^2+^ transients is of limited value. In our view, the Ca^+^ transients are best viewed as a binary measure: success or failure (Figure 4D). Future experiments may be able to address a portion of these quantitative shortcomings by measuring NE release using recently-developed NE fluorescent receptors such as GRAB_NE_.^42^

Does the return of startle-evoked Ca^+^ transients that we observe 3-7 weeks after DSP4 treatment truly reflect NE release from regrown NE axons? Certainly, our morphological findings show that NE axon regrowth is well underway at these time points (Figures 1, 2, S1, S2, S4). However, there is also the formal possibility that DSP4 inhibits NE release from survived axons for several weeks following treatment. If this is the case, the restoration of startle induced Ca^2+^ fluctuations in neocortical astrocytes may not entirely be a result of functional NE axon regrowth but rather may reflect, in part, the functional recovery of survived axons.

In our prior description of serotonin axon regrowth following amphetamine lesion, we employed serotonin cyclic voltammetry in the neocortex to assess serotonin release.^17^ This required stimulation of the medial forebrain bundle with an indwelling electrode at frequencies and durations greater than occur naturally.^43,44^ In those experiments we found that evoked serotonin release returned 6 months (but not 3 months) after the lesion of serotonin axons. Here, by examining NE release elicited by a natural physiological stimulus, we provide more definitive evidence supporting the functionality of these regrown monoaminergic axons.

Unlike neurons in the central nervous system, peripheral neurons are capable of axon regeneration following injury.^1,2,3^ Over the past 30 years, this field has been seeking to uncover the mechanisms underlying peripheral axon regeneration in the hope of using that knowledge to promote axon regeneration, and hence functional recovery, following CNS injury. This approach has been largely unsuccessful.^45^ Here, we characterized robust functional axon regrowth from neurons within the adult mammalian brain without experimental intervention such as gene deletion or tissue grafts.

When the present results are taken together with previous reports, it is now clear that both NE and serotonin axons can regrow following several forms of injury (including chemical lesions, neocortical stab and controlled cortical impact) in the adult rodent brain.^5,6,7,8,9,10,11,12,13,14,15,16,17,18,20,22,25,26,27^ Notably, axons of both types of monoaminergic neuros regrow in similar ways after chemical lesion. Both NE and serotonin axons are very stable in the absence of injury. Following toxic challenge, both regrow slowly and this regrowth is dominated by new axons entering the volume of view rather than sprouting from surviving axons. In both cases, the surviving axons continue to survive at normal rates, as do the regrown axons. Importantly, neither NE nor serotonin axons regrow using survived axons as a guide nor do either preferentially regrow along the paths previously occupied by lesioned axons (making it unlikely that they are following contact-mediated cues left behind when those lesioned axons regressed).

The similar dynamic properties of regrowing NE and serotonin axons suggests that these monoaminergic neurons may share a common molecular and genetic program for axon regrowth. In this vein, expression profiling of locus coeruleus NE neurons and raphe complex serotonin neurons together with various forms of injury may begin to reveal such a program and, in so doing, suggest pathways towards therapies to promote axon regeneration and functional recovery in the CNS.

## Materials and Methods

### Animals

All animal procedures were approved by the Johns Hopkins Medical Institute’s Institutional Animal Care and Use Committee. Both male and female mice were used for all experiments. Up to five mice were housed per cage under a 12-hour light/dark cycle, with food and water provided ad libitum. To selectively label NE neurons with EGFP for the initial evaluation of axon regrowth in fixed tissue (Figures 1, S1, and S2), Dopamine-β-hydroxylase (DBH)-cre mice (GENSAT, STOCK Tg(DBH-cre)KH212Gsat/Mmucd, Stock #032081-UCD, RRID: MMRRC_032081-UCD)^46^ were crossed with the mTmG reporter line (Jackson Laboratory, Stock #007576, Gt(ROSA)26Sortm4(ACTB-tdTomato,-EGFP)Luo/J, RRID: MGI:3722405).^34^ To selectively label NE neurons with TdTomato for the in vivo imaging experiments (Figures 2, 3, and S3-5), DBH-cre mice were crossed with the Ai14 reporter line (B6;129S6-Gt(ROSA)26Sortm14(CAG-tdTomato)Hze/J, Stock # 007908, RRID:IMSR_JAX:007908).^47^

### Surgical procedures for in vivo imaging

Mice, aged 9-12 weeks, received the following surgical procedures in preparation for in vivo imaging experiments. A small dose of dexamethasone (0.2ml at 0.04mg/ml) and lidocaine (0.1ml of 0.5%) were administered subcutaneously to the scalp prior to surgery to minimize potential swelling at the surgical site. Systemic Buprenex (0.1mg/kg at 0.006mg/ml) as an analgesic was given at the beginning of surgery. Surgical anesthesia was administered using isoflourane (5% induction, reduced to 1.5-2% during surgery). Following deep anesthesia, as assessed by paw pinch reflex and breathing rate, the mouse was placed in a stereotaxic device (Stoelting), and paralube was applied to the eyes and the skin overlying the skull was shaved. A cranial window (2.2 x 2.2 mm) was created on the skull above the somatosensory cortex (anterior-medial corner located −1mm posterior and +1mm lateral from bregma) with a hand drill (Fine Science Tools). The surface of the skull was periodically bathed in saline and drilling was intermittent to ensure that the underlying cortex was not damaged due to excessive heat from drill bit rotation. Once the skull flap was excised and the dura exposed, a glass pipet was filled with mineral oil and then connected to a Nanoject II device (Drummond Scientific Company) and front filled with AAV5-gfaABC1D-jGCaMP8s (pZac2.1-GfaABC1D-lck-jGCaMP8s, a gift from Prof. Loren Looger; Addgene plasmid # 176761 ; RRID:Addgene_176761) and viral prep was conducted by either the Janelia Viral Tools team or the Penn Vector Core. 40.2nl of virus was injected at a depth of 50µm and then again at a depth of 200um into two locations that bisected the anterior-to-posterior axis and trisected the medial-to-lateral axis of the exposed dura. These specific locations were chosen to avoid puncturing large surface blood vessels. Surgifoam (Johnson and Johnson) soaked in artificial cerebrospinal fluid (ACSF) was applied to the dura in order to stop any bleeding. Following injection, a square glass coverslip was placed inside the walls of the craniotomy and superglue was applied at the edges to form a cranial window, to cover the exposed skull, and to fasten head bars for in vivo imaging. Following this procedure, a subcutaneous injection of Baytril (2.5 mg/kg) was given as antibacterial prophylaxis.

### DSP4 and saline injection protocol

4-6 weeks following viral injection and cranial window installation surgery, mice received a single 50mg/kg intraperitoneal dose of DSP4 (N-(2-chloroethyl)-N-ethyl-2-bromobenzylamine, Tocris Bioscience #2958, PubChem ID: 38533)^24^ reconstituted in sterile water or an injection of volume-matched saline as a control.

### Immunohistochemistry

Mice were deeply anesthetized with ketamine (100mg/kg) and xylazine (10mg/kg) and intracardially perfused with ice-cold 4% paraformaldehyde (PFA) in phosphate buffer solution (PBS). Brains were removed and postfixed in 4% PFA at 4°C overnight before 48 hours of cryoprotection in graded sucrose steps (15% and 30% for 24 hours each). Whole brains were sectioned in the sagittal plane on a freezing sliding microtome (Leica) to a thickness of 40µm and stored at 4°C in PBS with sodium azide (Fischer, 0.002%) until use. Brain sections were then washed in 0.3% TritonX100 PBS and blocked for 2 hours at room temperature in 5% normal goat serum. Following blocking, sections were incubated at 4°C overnight in primary antibodies. The following primary antibodies were used: mouse anti-NET (Invitrogen #MA5-24647, RRID: AB_2637262, 1:200), chicken anti-GFP (Aves Labs #GFP-1010, RRID: AB_2307313, 1:6000), rabbit anti-DsRed (Takara Biosciences #632496, RRID: AB_10013483, 1:1000) and mouse anti-S100β (Sigma-Aldrich, HPA# AMAB91038, 1:000). Following primary incubation, the sections were washed in 0.3% TitonX100 PBS 3 times for 5 minutes each and then incubated with secondary antibodies. The following secondary antibodies were used: Alexa Fluor 488-labeled goat anti-mouse (Jackson ImmunoResearch Labs, catalog #115-545-003, RRID: AB_2338840, 1:500) Alexa Fluor 488-labeled goat anti-chicken (Jackson ImmunoResearch Labs, catalog #103-545-155, RRID:AB_2337390; 1:500), Alexa Fluor 594-labeled goat anti-mouse (Jackson ImmunoResearch Labs, catalog #115-586-146, RRID: AB_2338899; 1:500), Alexa Fluor 594-labeled goat anti-rabbit (Jackson ImmunoResearch Labs, catalog #111-585-144, RRID:AB_2307325; 1:500). Sections were again washed 3 times for 5 minutes each in 0.3% TritonX100 PBS, mounted on slides, and coverslipped using Prolong Antifade Diamond mounting media. Z-stack images were acquired with a 1µm step size using a laser scanning confocal microscope (Zeiss LSM 880 AxioExaminer.Z1, RRID:SCR_020925, 40x NA1.30 oil objective).

### In vivo imaging of NE axons

Mice were lightly anesthetized with 1%-2% isoflurane, placed on a feedback-controlled warming pad set to 37.5°C, and head fixed by fastening the lateral bars of the stereotaxic microscope stage to the mouse’s head bars. The stage was fixed on an x-y translator under a laser-scanning confocal microscope (Zeiss LSM 880 AxioExaminer.Z1, RRID:SCR_020925, Zeiss 20X NA 1.0 objective) and a non-descanned photomultiplier tube attached to the epifluorescence port. Two-photon excitation (1040 nm) was provided by a Coherent MRU X1 Laser producing ∼23-45 mW of power measured with a parked beam at the back aperture of the objective lens. Z-stacks were acquired starting at a depth of 150µm and continued in the dorsal direction to the cortical surface with a step size of 1µm. The laser power was consistently attenuated by software as the stack progressed from the starting depth to the cortical surface. Stacks were acquired every other week until the image windows clarity has deteriorated. A few time points were omitted when the microscope was being serviced and hence was unavailable.

### In vivo imaging of GCaMP8s

Awake mice were placed on a custom-designed rotating-disc treadmill and head fixed by fastening the lateral bars of the stereotaxic microscope stage to the mouse’s head bars. The treadmill was freely moving and placed under a Moveable Objective Microscope (Sutter Instruments) and two-photon excitation (920nm) was provided by a Spectra Physics Deep See infrared laser coupled to a Pockels cell and beam expanding telescope with fast resonant-galvo scanning. Time series image acquisition occurred at a rate of 30 frames/s. We averaged 6 frames per timepoint.

### Air puff induced startle responses

Mice were habituated to handling and head-fixation to the custom stereotaxic-treadmill microscope stage over the course of seven consecutive days prior to experimental recording. On days of data collection, head-fixed mice were allowed to rest on the treadmill for a 15-minute habituation period before the first trial was initiated. Each trial consisted of a total of 40 seconds. Time series image acquisition recorded baseline activity for the initial 10s before an automated delay triggered the application of an air-puff stimulus (compressed air applied at 30psi for 1s) to the back of the mouse’s neck. Time series image acquisition continued for another 30s to capture the response (Figure 4). The baseline recording period was limited to 10s to reduce phototoxicity. On experimental data collection days, mice were given 4 consecutive trials to account for the natural variation in the mouse’s state of vigilance which can trigger unstable baseline recordings or temporarily saturate the mouse’s response capacity. Trials were spaced 5 minutes apart to reduce habituation to the air-puff stimulus. For the α1-adrenoreceptor antagonist experiment (Figure 4 B-C), each animal underwent 3 consecutive trials before and after treatment with prazosin (7.5mg/kg, Tocris Biosciences, Cat. #0623, PubChem ID# 68546) separated by a 15-minute interval.

### Image analysis NE Axons

Fiji’s (RRID:SCR_002285) Labkit Pixel Classifier was used to isolate biologically relevant signals in both confocal (for immunohistochemistry) and in vivo 2-photon z-stacks. To calculate total surface area, z-stacks were reconstructed in Imaris (RRID:SCR_007370) and total surface area was determined using the automatic creation function within the Surface Creation Wizard. Exhaustive hand tracing of axon length was performed on a subset of images in a reference animal using Imaris to confirm that the relative change in total surface area between week 2 and week 8 following DSP4 treatment was similar to the relative change in total axon length (17% and 18.7% increase respectively). To classify axons as new, survived, or sprouted, Labkit-processed images of each data collection timepoint were manually reconstructed for each mouse into longitudinal time series in Imaris and aligned using 3-rigid body alignment based on common structural features. The aligned-image volumes for each mouse were then compressed into a single volume as separate channels. Axon surfaces were then generated and broken into individual segments using the seed-point generation function within Imaris. A Python-generated Center of Mass plugin was then used to create a spot at the center of mass of each surface segment. Segments were then classified as follows:

- Survived: 2 weeks following treatment, axon segments were classified as survived if their center of mass lay within 5µm of the center of mass of an axon segment present 1 day prior to treatment. In all subsequent weeks, survived axon segments were defined as those whose center of mass lay within 5µm of the center of mass of an axon segment classified as survived 2 weeks following treatment.
- New: 2 weeks following treatment, axon segments were classified as new if their center of mass lay 5µm or further from the center of mass of an axon segment present 1 day prior to treatment. In all subsequent weeks, new axon segments were defined as those whose center of mass lay 5µm or further from the center of mass of an axon segment classified as survived 2 weeks following treatment.
- Sprouted: sprouted axon segments were defined as elaborated growth directly from the trunk of survived axons, these segments were exhaustively identified and classified by eye.

### Startle-induced GCaMP response

Time series images for each trial were loaded into ImageJ/Fiji and a mean spatial smoothing filter of 1 was applied, averaging the intensity of each pixel with each of its neighbors. A background intensity threshold was determined and then a 75 x 75µm ROI was placed by eye over the location with the highest GCaMP fluorescent coverage. The total intensity of all pixels within that ROI which exceeded the background threshold for each frame were compiled and imported into R for further processing and analysis. ΔF/F values were calculated and a cubic smoothing spline (R Package: *npreg* version 1.0-9) was applied. Individual trials were excluded if the baseline range exceeded 0.5 ΔF/F. GCaMP responses were scored as successful if they met the following criteria: peak amplitude ΔF/F > 0.5 and a peak z-score > 6.

### Animal exclusion criteria

Animals were excluded from further analysis at several points during the experimental timeline. Animals were entirely excluded from the experiment at the outset due to inadequate clarity of the imaging window or insufficient expression of the AAV5-gfaABC1D-jGCaMP8s virus. Expression was deemed insufficient if less than 10% of pixels in the 75x75µm ROI failed to exceed the background intensity threshold. Animals were entirely excluded 1 week prior to treatment if they failed to demonstrate a GCaMP response to the air-puff startle stimulus in any of the 4 trials. Animals were excluded from data collection at subsequent time points due to deterioration of the imaging window clarity over time.

### Data Processing and Statistical analysis

Linear regression was used to compare numeric dependent variables. Generalized Linear model with Poisson distribution and logarithmic link was used to compare binary dependent variables. The models included treatment, time (in weeks) and treatment by time interactions as well as cluster-correlated variance estimates for animal to account for potential within-animal correlations.

To compare the overall trajectories for numeric measures in Figure 3, the area under the curve (AUC) was calculated using the trapezoid method. These AUCs were compared using Wilcoxon signed-rank test for paired data. Direct comparisons across treatment groups in Figure 1 and S4 were performed using unpaired student’s t test.

Statistical analyses were performed using STATA statistical software program StataCorp. 2023. Stata Statistical Software: Release 18. College Station, TX: StataCorp LLC. Data processing and student’s t tests were conducted using R Project for Statistical Computing (R) (RRID:SCR_001905).

## Acknowledgments

Julia Brill and Haley Janowitz provided guidance in the research and provided helpful comments on the manuscript. Statistical analysis was performed by Gayane Yenokyan of the Johns Hopkins Institute for Clinical and Translational Research. Aleksandr Smirnov and the Neuroscience Imaging Center (NIC) in the Solomon H. Snyder Department of Neuroscience provided instrumentation and support for in vivo 2-photon and confocal imaging.

## Author Contributions

P.C. designed, performed, and analyzed all experiments and co-wrote the manuscript. D.J.L directed the overall experimental design and analysis and co-wrote the manuscript.

## Supporting Information for

**Fig. S1.**
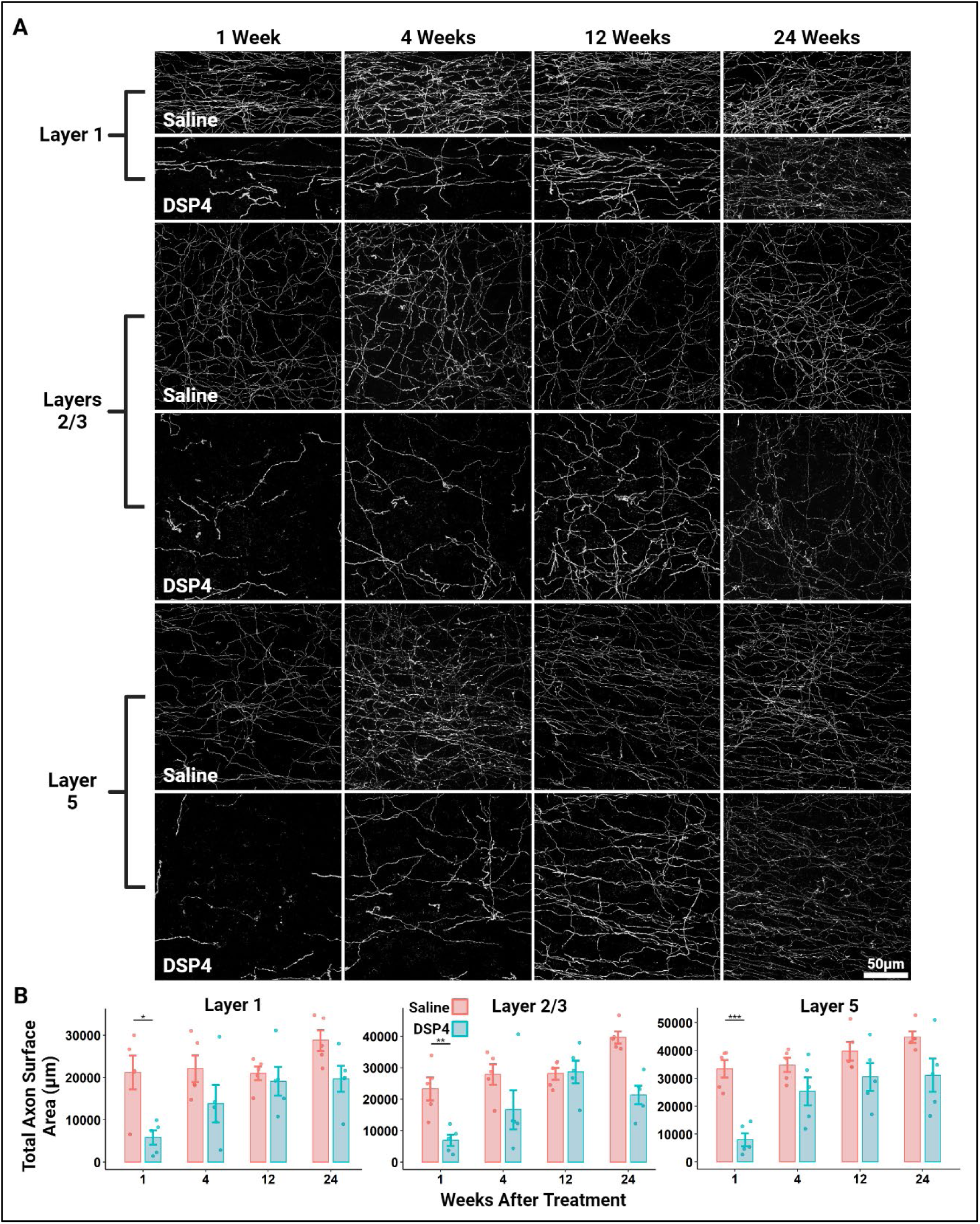
NE axons innervating the adult mouse primary somatosensory cortex regrow following a chemical lesion with the NE axon-specific neurotoxin DSP4. ***A***, Representative 30µm maximum projected confocal stack images of layers 1, 2/3, and 5 of the primary somatosensory cortex in mice sacrificed 1-, 4-, 12-, and 24-weeks following treatment with DSP4 (50mg/kg) or saline. Dopamine-β-hydroxylase (DBH)-cre x mTmG mice were used to selectively label NE axons, neuronal tissue was sliced along the sagittal plane, and the native signal was amplified through processing with antibodies raised against GFP. ***B***, IMARIS software was used to quantify the total symbol represents the total axon surface area of a single sagittal section of an individual mouse (n=5/group) and vertical bars show the standard error. * = *P* < 0.05; ** = *P* < 0.01; *** = *P* < 0.001.

**Fig. S2.**
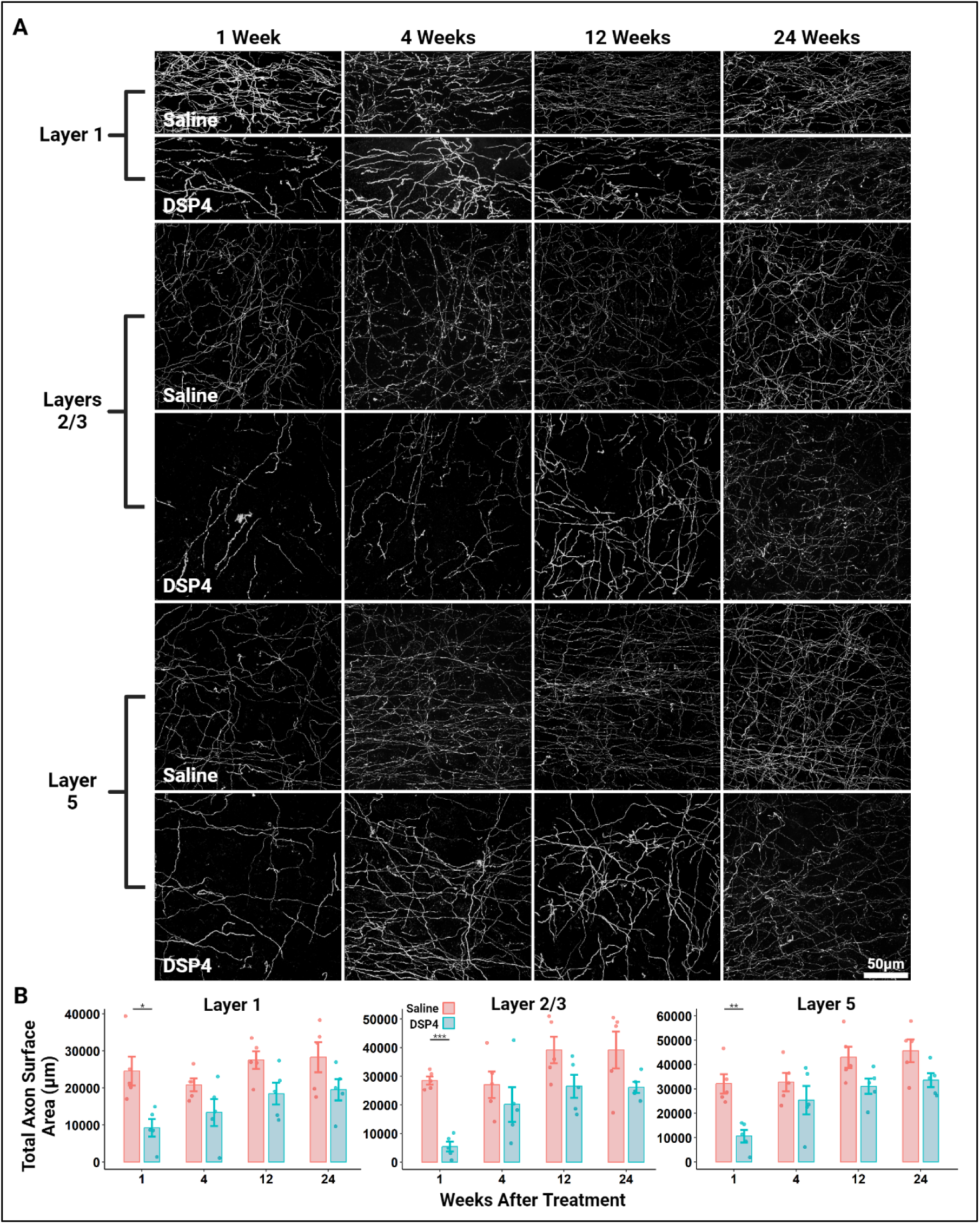
NE axons innervating the adult mouse primary motor cortex regrow following a chemical lesion with the NE axon-specific neurotoxin DSP4. ***A***, Representative 30µm maximum projected confocal stack images of layers 1, 2/3, and 5 of the primary motor cortex of mice sacrificed 1-, 4-, 12-, and 24-weeks following treatment with DSP4 (50mg/kg) or saline. Dopamine-β-hydroxylase (DBH)-cre x mTmG mice were used to selectively label NE axons, neuronal tissue was sliced along the sagittal plane, and the native signal was amplified through processing with antibodies raised against GFP. ***B***, IMARIS software was used to quantify the total axon surface area within demonstrated in *A*. Each plot symbol represents the total axon surface area of a single sagittal section of an individual mouse (n=5/group) and the vertical bars show the standard error. * = *P* < 0.05; ** = *P* < 0.01; *** = *P* < 0.001.

**Fig. S3.**
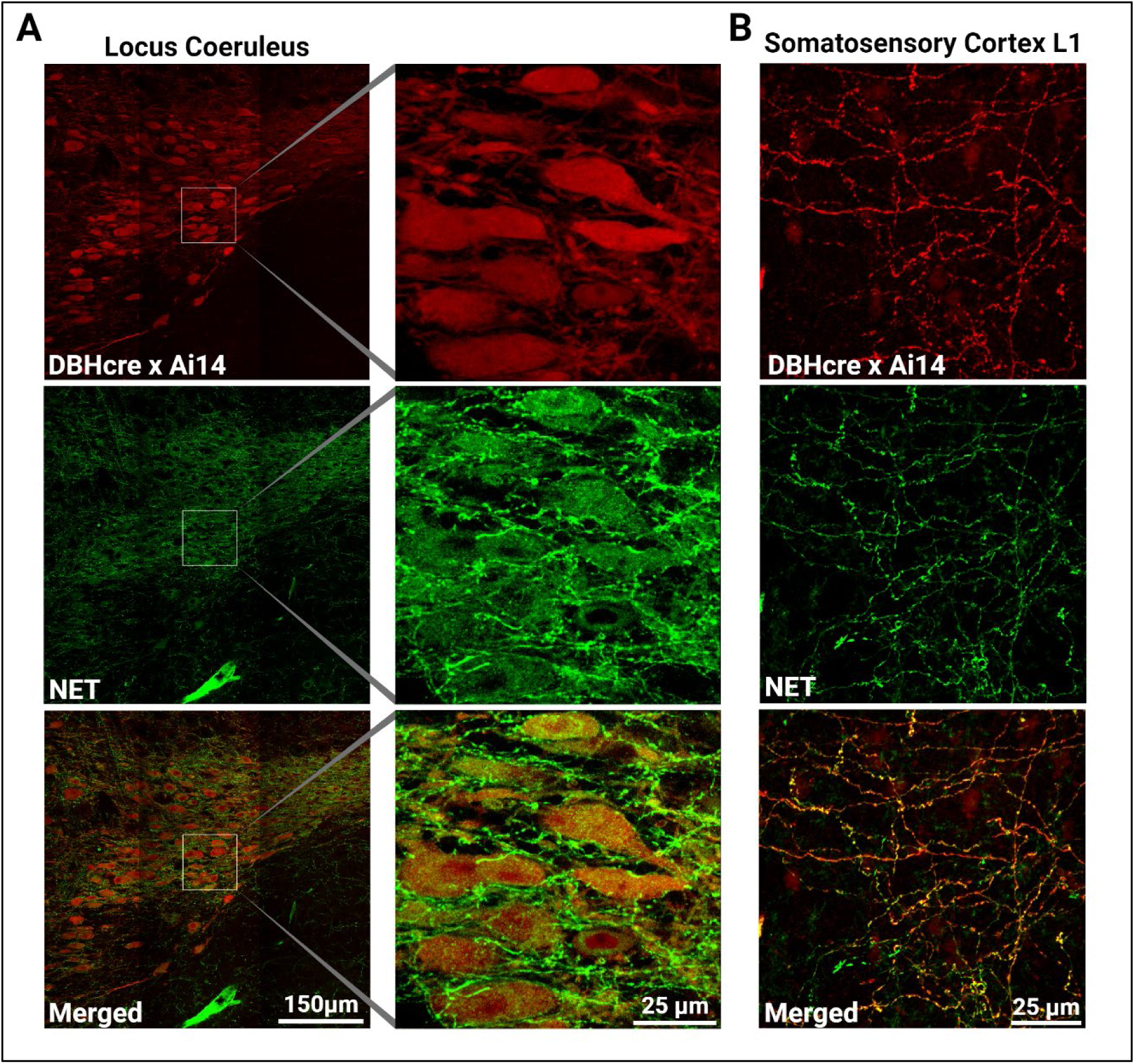
NE neurons are selectively labeled with TdTomato in dopamine-β-hydroxylase (DBH)-cre x Ai14 transgenic mice. ***A***, Representative 3-by-3 tiled (left) and magnified subsection (right) 30µm maximum projected confocal stack images of the locus coeruleus. Brain tissue was sliced in the sagittal plane and counterstained with antibodies raised against the norepinephrine transporter (NET) to selectively label the plasma membrane of NE neurons. Ai14-driven tdTomato fluorescence was amplified with cross-reactive antibodies raised against DsRed. Ai14-positive cell bodies and axons consistently show NET immunoreactivity. ***B***, Representative 30µm thick maximum projected confocal stack images of layer 1 of the primary somatosensory cortex section were processed in the same manner as those in panel *A.* Ai14-positive axons in this region consistently show NET immunoreactivity as well.

**Fig. S4.**
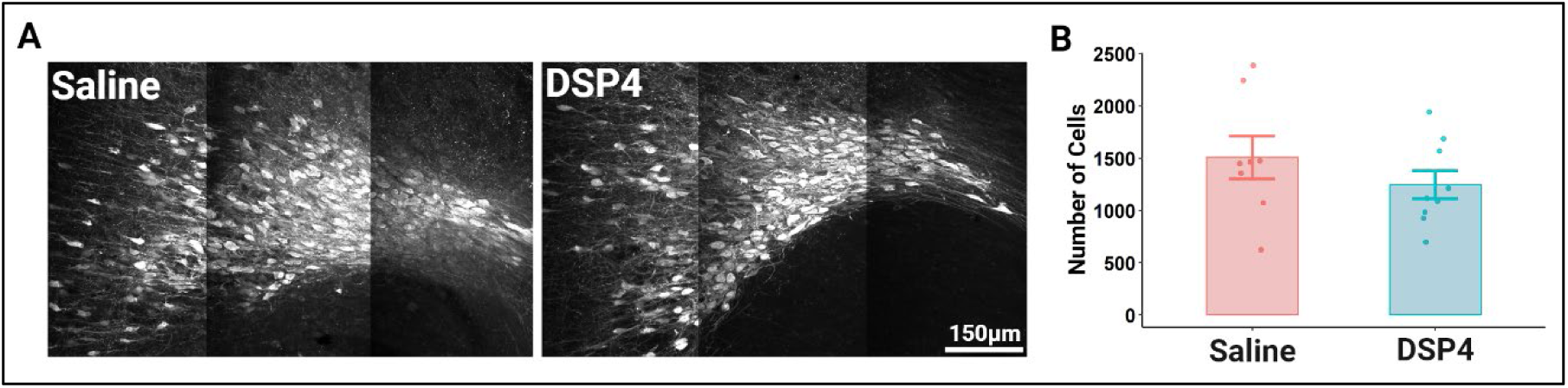
A single dose of DSP4 (50mg/kg), as used throughout the present study, does not lesion NE cell bodies within the locus coeruleus of the adult mouse. ***A***, Representative 3-by-3 tiled 30µm maximum projected confocal stack images of the locus coeruleus 24-weeks following treatment with DSP4 (50mg/kg) or saline. DBHcre x Ai14 mice were used to selectively label NE neurons with tdTomato. The brain was sliced along the sagittal plane, and the tdTomato signal was amplified using cross-reactive antibodies raised against DsRed. ***B***, IMARIS software was used to produce an exhaustive count of the Ai14-positive cell bodies within the locus coeruleus of individual animals. No significant difference in the number of cells was detected between the saline (n=8) and DSP4 (n=9) groups (p = 0.289). The standard error is represented by vertical bars.

**Fig. S5.**
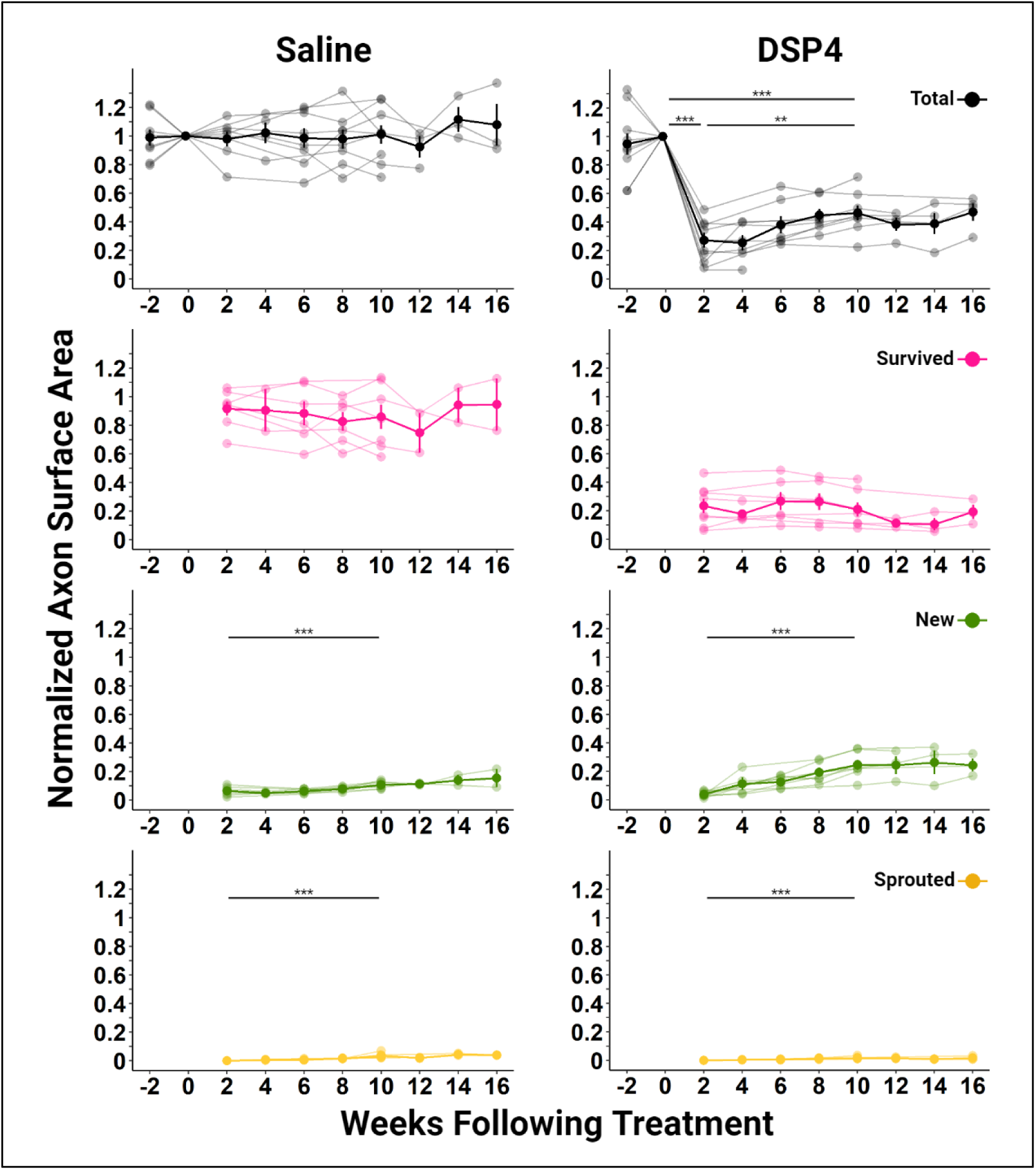
Long-term *in vivo* 2-photon imaging shows the regrowth of NE axons innervating the adult mouse somatosensory cortex following DSP4 treatment. Individual mouse (shaded) and population mean +/- standard error (solid) measurements of the axon surface area are shown. The population data are the same as illustrated in Figure 2B and are reproduced here to allow for comparison with the individual mouse measurements. Weeks during which data was able to be collected are indicated by shaded closed circles. Collection was sometimes limited by repair of the imaging rig and lines representing individual mice sometimes terminate due to deterioration of the optical quality of the imaging window. ** = *P* < 0.01; *** = *P* < 0.001.

**Fig. S6.**
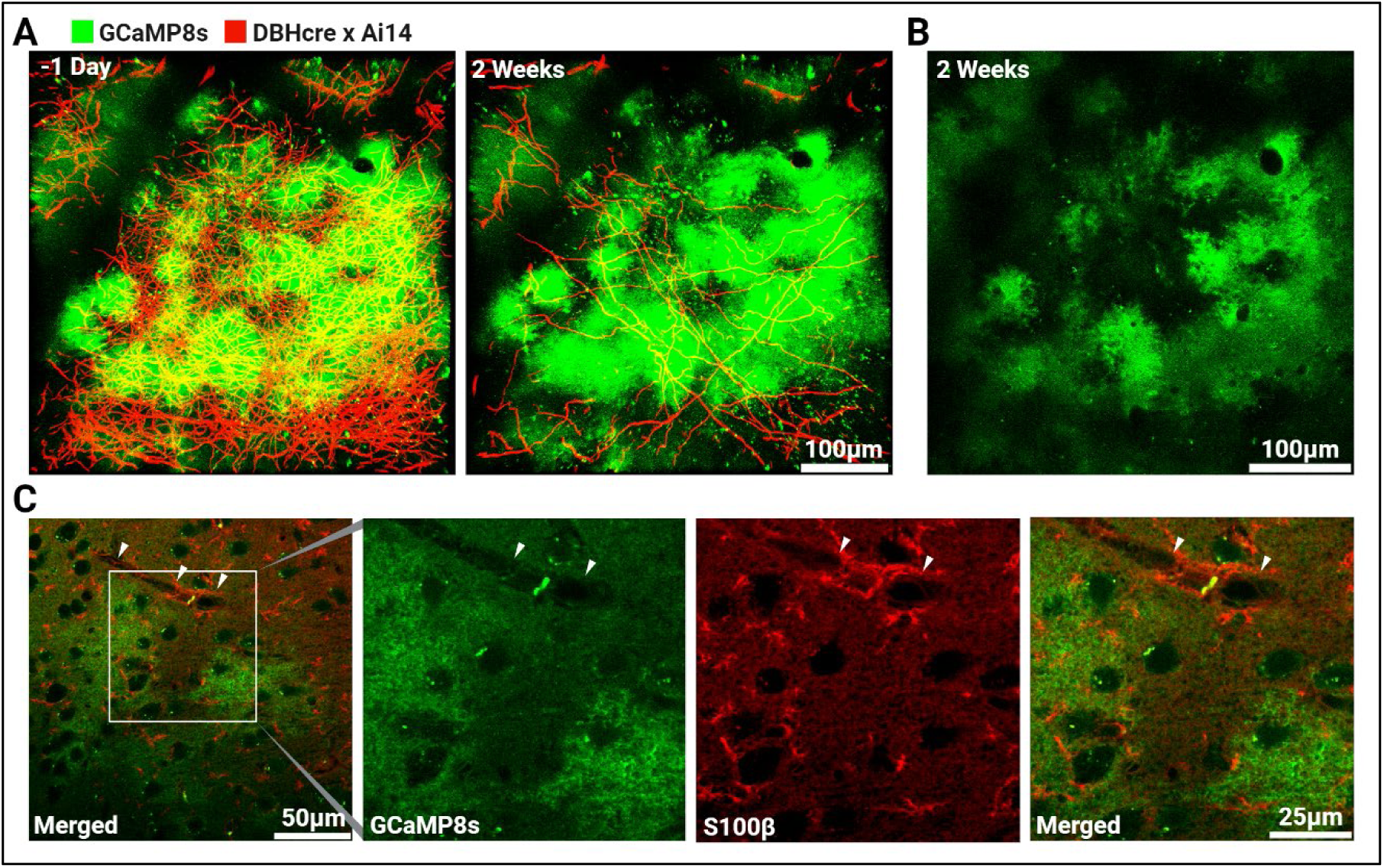
Infection with AAV5-gfaABC1D-jGCaMP8s induces expression of GCaMP8s in cortical astrocytes. ***A***, Representative 108µm 3-D reconstructed images of the primary somatosensory cortex show virally induced GCaMP8s (green) and DBHcre x Ai14 (red) one day prior to and two weeks following lesion with the NE-specific neurotoxin DSP4. Viral injection 7 weeks prior to lesion induces patchy GCaMP expression which shows the tiled patterning characteristic of astrocytes. Note that the expression pattern and intensity of GCaMP was not significantly altered over the 15 day-long span encompassing the DSP4 challenge. ***B***, 3µm-thick maximally projected z-stack images of the same GCaMP8s signal shown in *A* reveals the tendril-like processes characteristic of astrocytes. ***C***, Representative single image of layer 2/3 of the primary somatosensory cortex 16 weeks following infection with AAV5-gfaABC1D-jGCaMP8s to selectively induce GCaMP8s in cortical astrocytes. Fixed brain tissue was sliced in the sagittal plane and processed with antibodies raised against the astrocyte marker S100β. As in *A* and *B,* the GCaMP8s signal demonstrates a patchy transfection pattern and tiling while S100β labels all astrocytes, highlighting particular cytoskeletal elements. White arrowheads indicate a capillary contacted by S100β-positive astrocyte endfeet.

